# A mouse SWATH-MS reference spectral library enables deconvolution of species-specific proteomic alterations in human tumour xenografts

**DOI:** 10.1101/2020.02.03.930248

**Authors:** Lukas Krasny, Philip Bland, Jessica Burns, Nadia Carvalho Lima, Peter T. Harrison, Laura Pacini, Mark L. Elms, Jian Ning, Victor Garcia Martinez, Yi-Ru Yu, Sophie E. Acton, Ping-Chih Ho, Fernando Calvo, Amanda Swain, Beatrice A. Howard, Rachael C. Natrajan, Paul H. Huang

**Author notes:** Correspondence to: Paul H Huang, Division of Molecular Pathology, Institute of Cancer Research, 237 Fulham Road, London SW3 6JB, United Kingdom.

## Abstract

SWATH-mass spectrometry (MS) enables accurate and reproducible proteomic profiling in multiple model organisms including the mouse. Here we present a comprehensive mouse reference spectral library (MouseRefSWATH) that permits quantification of up to 10,597 proteins (62.2% of the mouse proteome) by SWATH-MS. We exploit MouseRefSWATH to develop an analytical pipeline for species-specific deconvolution of proteomic alterations in human tumour xenografts (XenoSWATH). This method overcomes the challenge of high sequence similarity between mouse and human proteins, facilitating the study of host microenvironment-tumour interactions from ‘bulk tumour’ measurements. We apply the XenoSWATH pipeline to characterise an intraductal xenograft model of breast ductal carcinoma in-situ and uncover complex regulation of cell migration pathways that are not restricted to tumour cells but also operate in the mouse stroma upon progression to invasive disease. MouseRefSWATH and XenoSWATH opens new opportunities for in-depth and reproducible proteomic assessment to address wide-ranging biological questions involving this important model organism.

## Introduction

Mass spectrometry (MS) has become an essential tool for contemporary proteomic research in life sciences. Conventional data-dependent acquisition (DDA) mode, where a fixed number of the most abundant precursor ions in survey scans are automatically selected for fragmentation, enables the identification of thousands of proteins in a single MS experiment. However, the stochastic nature of precursor ion selection in DDA leads to low reproducibility in peptide identification between experimental runs [1, 2]. Sequential window acquisition of all theoretical mass spectra (SWATH-MS) or data-independent acquisition mass spectrometry (DIA-MS) is a next-generation label-free quantification method that enables highly reproducible peptide identification and more accurate quantification in large-scale proteomic analyses across multiple experiments [3–5]. This method is based on the periodic fragmentation of all precursor ions in wide and adjacent isolation windows that cover the entire mass to charge ratio (m/z) range measured [3]. In this manner, all precursor ions generated from the sample under investigation are fragmented and their fragmentation spectra acquired for subsequent *in silico* analysis. These quantitative digital proteome maps have been shown to have wide applications in medical research such as the discovery of potential biomarkers and therapeutic targets [6–10] as well as tumour proteotyping for cancer stratification and classification [11, 12].

Key to the success of SWATH-MS is the use of retention-time calibrated spectral libraries to identify and quantify peptides from the complex fragment ion mass spectra encoded within digital proteome maps [13]. While most published SWATH-MS studies have utilised study-specific experimentally derived spectral libraries, this approach is time consuming and is prone to wide variation between laboratories due to the lack of standardisation in DDA data acquisition and library generation. Such variation results in study-specific bias and poor inter-laboratory reproducibility. The recent development of the algorithms that control for false discovery rate (FDR) in SWATH-MS datasets has opened up the possibility of building large reference spectral libraries that can be readily shared by the community, increasing data reproducibility across laboratories and accelerating SWATH-MS experiments without the need to generate study-specific spectral libraries [14, 15].

Several such large reference spectral libraries have been built and deposited into repositories such as SWATHAtlas (www.SWATHatlas.org) including libraries for human [15], fruit fly [16], zebrafish [17] and yeast [18]. There is however currently no publically available comprehensive mouse reference spectral library. The mouse as a model organism has and continues to be extensively used in developmental biology and medical research due to the complex genetics and physiological systems that mammals share. The array of innovative genetic strategies available for engineering mice has revolutionised our understanding of multiple human diseases including cancer, diabetes, autoimmune disease and heart disease amongst others. Furthermore, immunocompromised mouse models have been used for more than a decade as hosts for human tumour (both cell line and patient-derived) xenografts which have served as an essential tool to investigate the factors that drive carcinogenesis as well as to evaluate cancer therapeutics [19]. Developing a comprehensive mouse reference spectral library will facilitate the application of SWATH-MS to address key research questions involving this widely used model organism. A number of study-specific mouse spectral libraries have previously been reported [20–23], for instance a spectral library that is comprised of the mouse immunopeptidome containing 1573 peptides presented by MHC class I molecules [20] and a spectral library reported by Williams *et al*. which is comprised of 5152 proteins (30% of mouse proteome) generated from five organs [21]. Notably, in the aforementioned human tumour xenograft models, there is currently no established method to deconvolute host (mouse) versus tumour (human) proteomic alterations from ‘bulk tumour’ proteomes which has limited our ability to study the role of the tumour microenvironment in driving tumour initiation and progression.

In this study, we present a comprehensive mouse reference spectral library (MouseRefSWATH) generated from 15 distinct mouse organs and cellular samples. MouseRefSWATH was built from 254 individual MS experiments and is composed of transitions for 167,138 proteotypic peptides from 10,597 proteins representing 62.2% of manually validated mouse protein-encoding genes (Swissprot database). The performance of MouseRefSWATH was evaluated in two publicly available SWATH-MS datasets which showed both qualitative and quantitative reproducibility when compared to published study-specific spectral libraries. We further report the development of a novel application of MouseRefSWATH for SWATH-MS-based mapping of species-specific temporal proteomic alterations in an orthotopic tumour xenograft model of breast ductal carcinoma *in situ* (DCIS). Utilising this approach, we reveal for the first time, simultaneous temporal regulation of cell migration pathways in both the host mouse mammary gland and human tumour cells during the course of DCIS to invasive breast cancer (IBC) progression. This XenoSWATH pipeline for species-specific deconvolution of ‘bulk tumour’ proteomics data provides a useful tool with broad applications for the analysis of host microenvironment-tumour interactions in xenograft models without the need for prior separation of host and tumour cell populations. The MS raw data and MouseRefSWATH spectral library for this study is available via ProteomeXchange with identifier PXD017209.

## Results

### Proteomic analysis of murine tissue and cells

To maximize coverage of the mouse proteome in the reference spectral library, we performed DDA proteomic analysis of a comprehensive range of 7 murine organs comprising heart, brain, lung, liver, kidney, lymph node and mammary gland (Figure 1 and Table 1). In addition, we undertook proteomic profiling from primary CD8^+^ T-lymphocytes (non-stimulated and activated) as well as immortalised murine cell lines including normal (NF1) and cancer-associated (CAF1) fibroblasts [24] and commercially available cell lines NIH-3T3, C2C12, 4T1 and Ba/F3 of diverse tissue origin (Table 1). As outlined in the workflow shown in Figure 1, extracted proteins from each sample type were digested by trypsin and the resulting peptide mixture was subjected to offline fractionation in the first dimension either by strong cation exchange (SCX) chromatography or reverse phase chromatography in high pH (HpH-RP) (Table 2). To calibrate the chromatographic retention time for individual DDA runs, prior to liquid chromatography tandem mass spectrometry (LC-MS/MS) analysis each fraction was spiked with the iRT calibration standard that contains mixture of 11 synthetic peptides [13, 25]. Peptides from each fraction were subsequently analysed by LC-MS/MS in DDA mode and the acquired data were processed by SpectroMine [26, 27]. An average of between 3200-6600 proteins were identified across the different samples as shown in Table 2.

**Figure 1.**
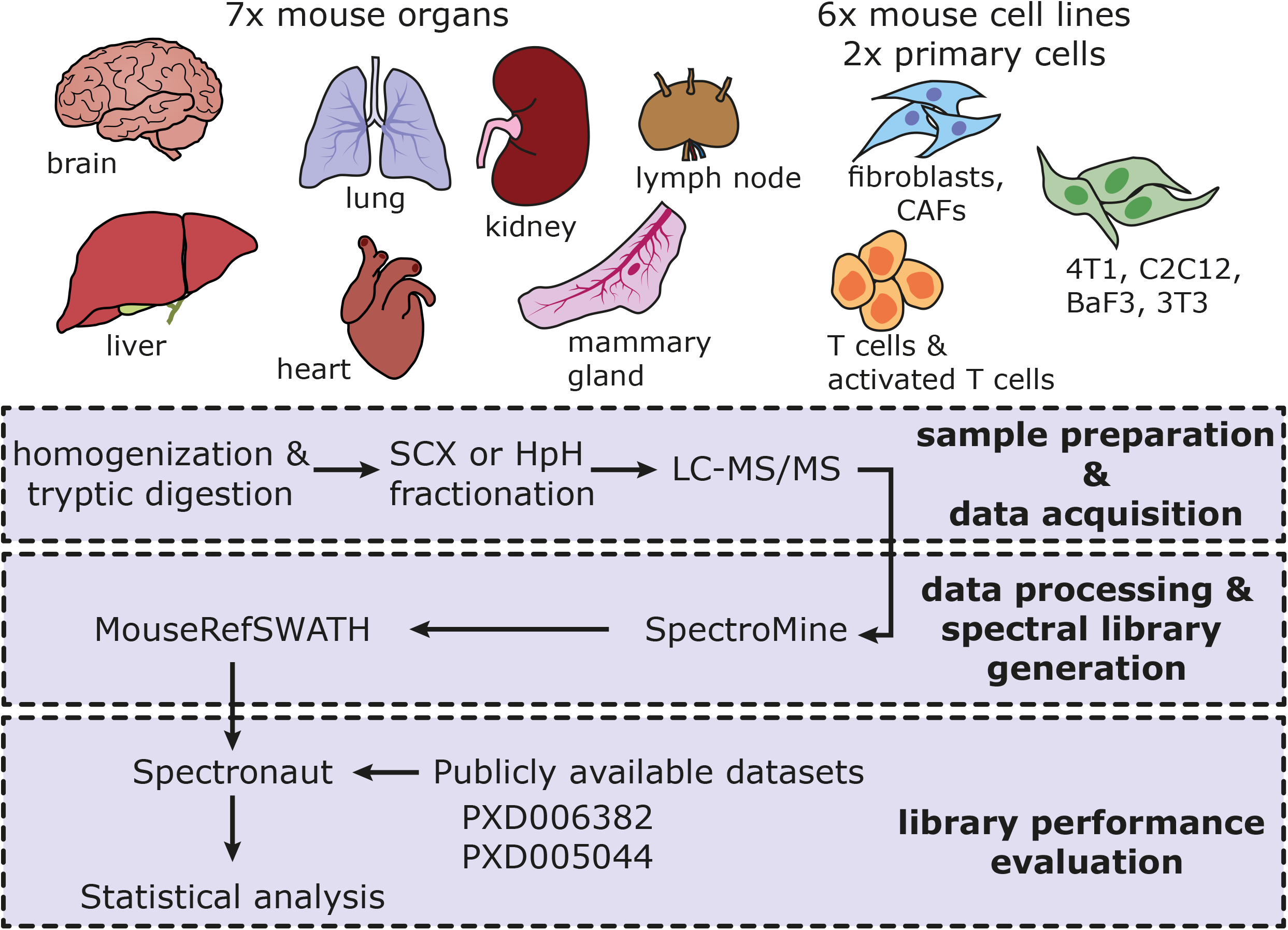
Overview of the sample types and schematic workflow used to build the MouseRefSWATH reference spectral library. 15 unique sample types comprising 7 mouse organs, 2 primary cell types and 6 mouse cell lines were analysed to maximise proteome coverage. The workflow is comprised of a first step of sample preparation and data acquisition of each sample by DDA-based LC-MS/MS. In the second step, the resulting proteomic data was subjected to processing and spectral library generation using the combined Search Archives approach in the SpectroMine software. The subsequent evaluation of the performance of the MouseRefSWATH library was undertaken with the Spectronaut software utilising two publicly available datasets.

**Table 1:** Overview of the samples used for mouse reference spectra library generation

| **Sample** | **Sample type** | **Mouse/cell strain** | **Detailed sample description** |
| --- | --- | --- | --- |
| **Liver** | tissue | SCID-beige mice | dissected from 14-18 week old female mouse |
| **Lung** | tissue | SCID-beige mice | dissected from 14-18 week old female mouse |
| **Mammary gland** | tissue | SCID-beige mice | dissected from 14-18 week old female mouse |
| **Brain** | tissue | NCR nude mice | dissected from 6 months old female mouse |
| **Kidney** | tissue | NCR nude mice | dissected from 6 months old female mouse |
| **Heart** | tissue | NCR nude mice | dissected from 6 months old female mouse |
| **Axillary and bronchial lymph nodes** | tissue | C57BL/6N mice | dissected from 2-4 months old mouse, males and females |
| **NIH/3T3** | cell line | ATCC CCL 92 | mouse embryonic fibroblast |
| **4T1** | cell line | ATCC CRL 2539 | mouse epithelial cell, mammary gland tumour origin |
| **Ba/F3** | cell line | ATCC 300 | mouse IL-3 dependent pro-B cell line |
| **C2C12** | cell line | ATCC CRL 1772 | mouse C3H muscle myoblast |
| **Cancer associated fibroblast (CAF1)** | cell line | CAF1 | immortalized fibroblast isolated from mammary carcinoma tissue of FVB/n mouse |
| **Normal fibroblast (NF1)** | cell line | NF1 | immortalized fibroblast isolated from normal mammary glands of FVB/n mouse |
| **T cells** | primary cells | OT-I | Primary CD8+ effector T cells isolated from splenocytes of OT-I mouse |
| **Activated T cells** | primary cells | OT-I | Primary CD8+ effector T cells isolated from splenocytes of OT-I mouse and stimulated by ovalbumin peptides |

**Table 2:** Overview of the DDA datasets used for the generation of the MouseRefSWATH reference spectral library. Protein and peptide FDR threshold of 1% was used on each individual dataset.

| Sample | Sample type | Fractionation | Number of runs | Total number of protein IDs* |
| --- | --- | --- | --- | --- |
| Liver | tissue | SCX | 12 | 4471 |
| Lung | tissue | SCX | 24 | 6295 |
| Brain | tissue | SCX | 24 | 6366 |
| Kidney | tissue | SCX | 12 | 5093 |
| Heart | tissue | SCX | 24 | 3284 |
| Axillary lymph node | tissue | SCX | 12 | 5643 |
| Bronchial lymph node | tissue | SCX | 12 | 6358 |
| Mammary gland | tissue | SCX | 24 | 5433 |
| 3T3 | cell line | SCX, HpH-RP | 24 | 5799 |
| 4T1 | cell line | SCX, HpH-RP | 20 | 6293 |
| Ba/F3 | cell line | HpH-RP | 8 | 5353 |
| C2C12 | cell line | SCX, HpH-RP | 20 | 6681 |
| CAF1 | cell line | SCX | 12 | 5390 |
| NF1 | cell line | SCX | 12 | 3702 |
| Normal T-cells | primary cells | SCX | 12 | 6060 |
| Activated T-cells | primary cells | SCX | 12 | 6018 |

### Building the MouseRefSWATH spectral library

We built the mouse reference spectral library (MouseRefSWATH) from the datasets listed in Table 2 by employing the SpectroMine software (Figure 1). Only unique protein-specific (proteotypic) peptides were used to generate the library which comprise 10,597 proteins (Figure 2A), representing 62.2% coverage of manually annotated mouse protein-coding genes (Swissprot, 26/10/2018). Comparative analysis demonstrates a superior proteome coverage versus published spectral libraries of other higher eukaryotic organisms (human – 51% [15] and zebrafish – 40.4% [17]) (Figure 2B). Figure 2C shows the distribution of unique peptides per protein group with 90.6% of proteins represented by >1 unique peptide in the spectral library and 46.7% of proteins being represented by >10 unique peptides. The contribution plot (Figure 2D) shows that 1949 proteins (18.4%) in the MouseRefSWATH library were detected in all analysed sample types used in the generation of the library. The brain contributed the highest number of proteins (551, 5.2%) to the MouseRefSWATH library, followed by T cells (165, 1.6%) and lung (127, 1.2%).

**Figure 2.**
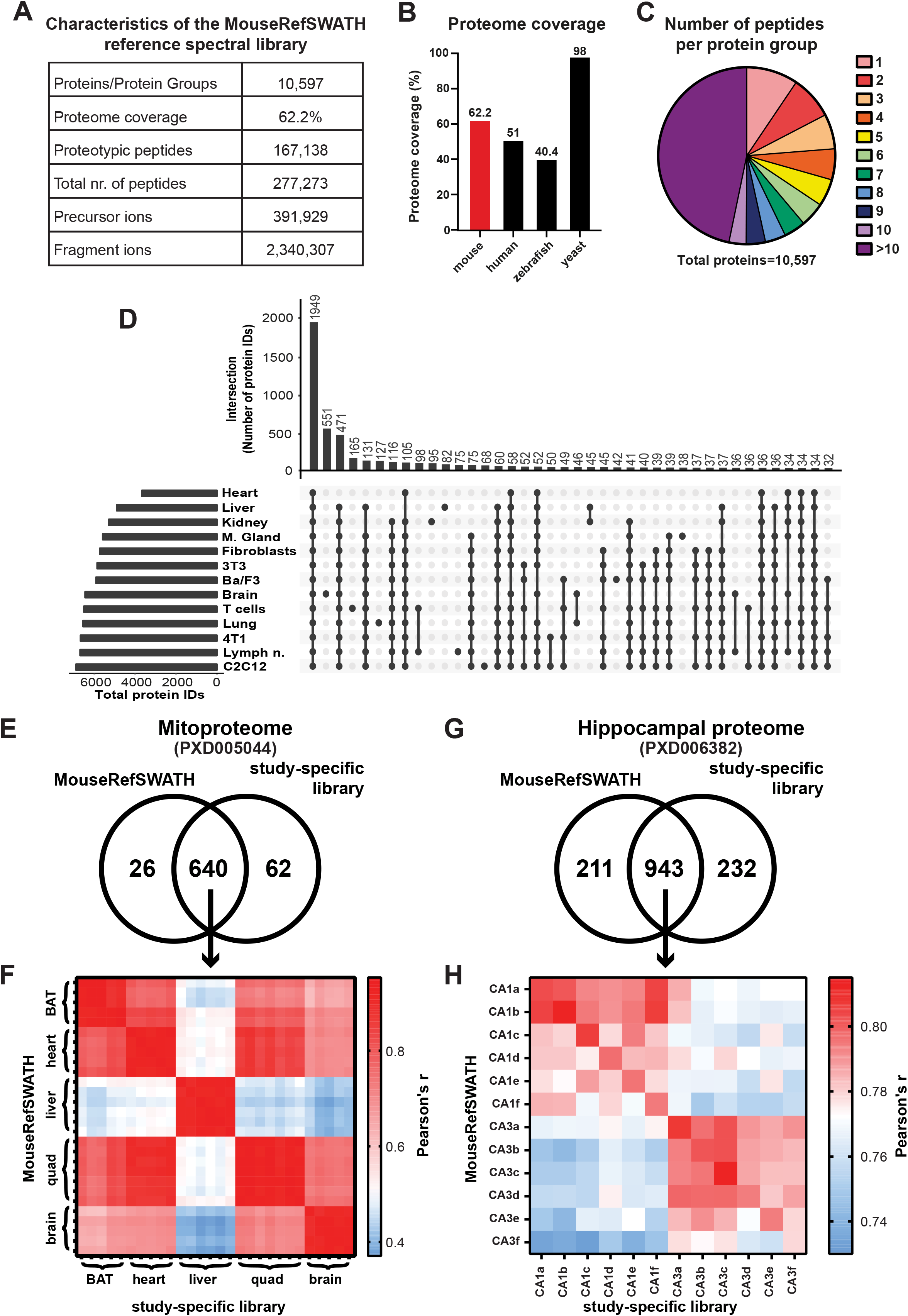
Characteristics and performance of the MouseRefSWATH reference spectral library. A) Detailed characteristics of the generated MouseRefSWATH library. B) Percentage proteome coverage of MouseRefSWATH library in comparison with published spectral libraries for other eukaryotic organisms, including human [15], zebrafish [17] and yeast [18]. C) Peptide coverage of the individual proteins in the MouseRefSWATH library. 90.6% of the proteins are represented by more than one unique peptide. D) Contribution plot of the tissue- or cell-specific proteins to the overall composition of the MouseRefSWATH library. E) Venn diagram depicting overlap of mitochondrial proteins detected by the MouseRefSWATH library and study-specific library based on analysis of the PXD005044 dataset [21]. F) Similarity matrix showing Pearson’s correlation coefficient of the 640 overlapping mitochondrial proteins which were quantified by either the MouseRefSWATH library or the study-specific library. 5-8 animals were used for each tissue site. BAT is brown adipocyte tissue and quad is quadriceps. G) Venn diagram depicting overlap of hippocampal proteins detected by the MouseRefSWATH library and study-specific library based on analysis of the PXD006382 dataset [22]. H) Similarity matrix showing Pearson’s correlation coefficient of the 943 overlapping hippocampal proteins which were quantified by either the MouseRefSWATH library or the study-specific library. Two hippocampal areas (CA1 and CA3) from 6 animals (a-f) were used.

### Evaluating the performance of the MouseRefSWATH spectral library

To demonstrate the utility of the MouseRefSWATH library and benchmark its performance against study-specific libraries generated as part of published SWATH-MS studies, we applied the reference spectral library to 2 publicly available SWATH-MS datasets focused on mitochondrial [21] and hippocampal [22] proteins (Figure 1). Each dataset was analysed using either the original study-specific library generated by the authors or the MouseRefSWATH library.

In a recent mitoproteome study undertaken by Williams *et al*. [21], a study-specific library based on DDA analysis of both total tissue lysates and mitochondria-enriched samples was utilised. We sought to assess the extent to which the MouseRefSWATH library was able to both qualitatively and quantitatively recapitulate the data generated by the study-specific library in the published study. The SWATH-MS data of mitochondrial proteins was downloaded from ProteomeXchange (PXD005044) and analysed using both libraries (Figure 2E & F). The Williams *et al*. experimental dataset was generated from mitochondria enriched from 5 distinct murine tissues (brown adipose tissue (BAT), heart, liver, quadriceps, brain,) from 5-8 animals. Qualitatively, the MouseRefSWATH library identified 91.1% (640 proteins) of the mitoproteome (Figure 2E) found using the study-specific library with an additional 26 proteins identified specifically by the MouseRefSWATH library. An assessment of the quantification of the 640 overlapping proteins showed a good correlation between both libraries with Pearson’s correlation coefficient r values ranging between 0.92 – 0.95 (Figure 2F).

We undertook a similar comparative analysis on the study by von Ziegler *et al.* [22] where the basal mouse hippocampal proteome was analysed utilising a study-specific spectral library built from DDA data generated from the mouse hippocampus. For this analysis, the SWATH-MS experimental data of hippocampal area CA1 and CA3 from 6 different animals were analysed (Figure 2G & H). The MouseRefSWATH library identified 80.2% (943 proteins) of hippocampus proteome found using the study-specific spectral library (Figure 2G). The MouseRefSWATH library further identified 211 more proteins compared to the hippocampus-specific spectral library. Good correlation in protein quantification between the MouseRefSWATH and study-specific library for the overlapping 943 proteins was observed with Pearson’s correlation coefficient r values ranging between 0.78-0.81 across the 6 mice in the experiment (Figure 2H). Taken together, our analysis demonstrates that applying the MouseRefSWATH to SWATH-MS datasets is routinely able to identify >80% of proteins identified using study-specific libraries with comparable quantification, highlighting its broad utility as a general reference library applicable for use in multiple mouse SWATH-MS datasets without the need to generate study-specific libraries.

### Developing the XenoSWATH analysis pipeline for deconvolution of mouse and human proteins in tumour xenograft SWATH-MS data

Mouse xenograft studies are one of the cornerstones of modern cancer research where human tumour cells are typically grafted into a mouse host either subcutaneously or orthotopically as a means to evaluate oncogene and tumour suppressor gene function or investigate the therapeutic effects of drugs [28, 29]. In these models, there are complex interactions between the human tumour cells and the host microenvironment which play important roles in driving cancer progression and therapy response [30–32]. There is currently no means to distinguish proteins of human and mouse origin from ‘bulk tumour’ proteomics data generated by SWATH-MS, which presents a key barrier in our ability to undertake deep characterisation of mechanisms of tumour-microenvironment interactions *in vivo*. Previous studies have attempted to tackle this challenge by separating human tumour and murine host cells utilising immunoaffinity cell enrichment strategies prior to MS analysis [33]. However, such methods are labour intensive, introduce sample preparation biases and disrupt the *in situ* architecture and cellular interactions important for driving tumour biology.

To address this challenge, we developed a novel pipeline (XenoSWATH) to deconvolute species-specific proteomic profiles from SWATH-MS data obtained in mouse xenograft experiments. In this XenoSWATH workflow (Figure 3), both the MouseRefSWATH library as well as a previously published pan-Human reference library [15] were used in the Spectronaut software [14]. In the first step of data processing, peptides were identified by searching the acquired SWATH-MS data against either the MouseRefSWATH or pan-Human library. Both libraries contain proteotypic peptides which can distinguish individual proteins, however given that humans and mice share ~70% of protein coding sequences [34], not all of these proteotypic peptides are also species-discriminating. Therefore to selectively quantify human and mouse proteins from ‘bulk tumour xenograft’ proteomic datasets, we focused on peptides that are both protein- and species-discriminating (see Figure 3 for example). To achieve this, rather than using individual FASTA files comprising *in silico* digested peptides for either human or mouse proteins, we instead manually combined both files into a single FASTA file for peptide quantification. This modification to the data processing pipeline enables Spectronaut to compare sequences of the identified peptides from either the mouse or human reference spectral library with the sequences of the *in silico* digested peptides within the combined FASTA file; and filter out any peptides that are shared between human and mouse. This leads to the retention of only peptides that are both proteotypic and species-discriminating for peptide quantification. The resultant output from this XenoSWATH pipeline is two datasets - one comprising entirely of murine proteins and the other of human proteins.

**Figure 3.**
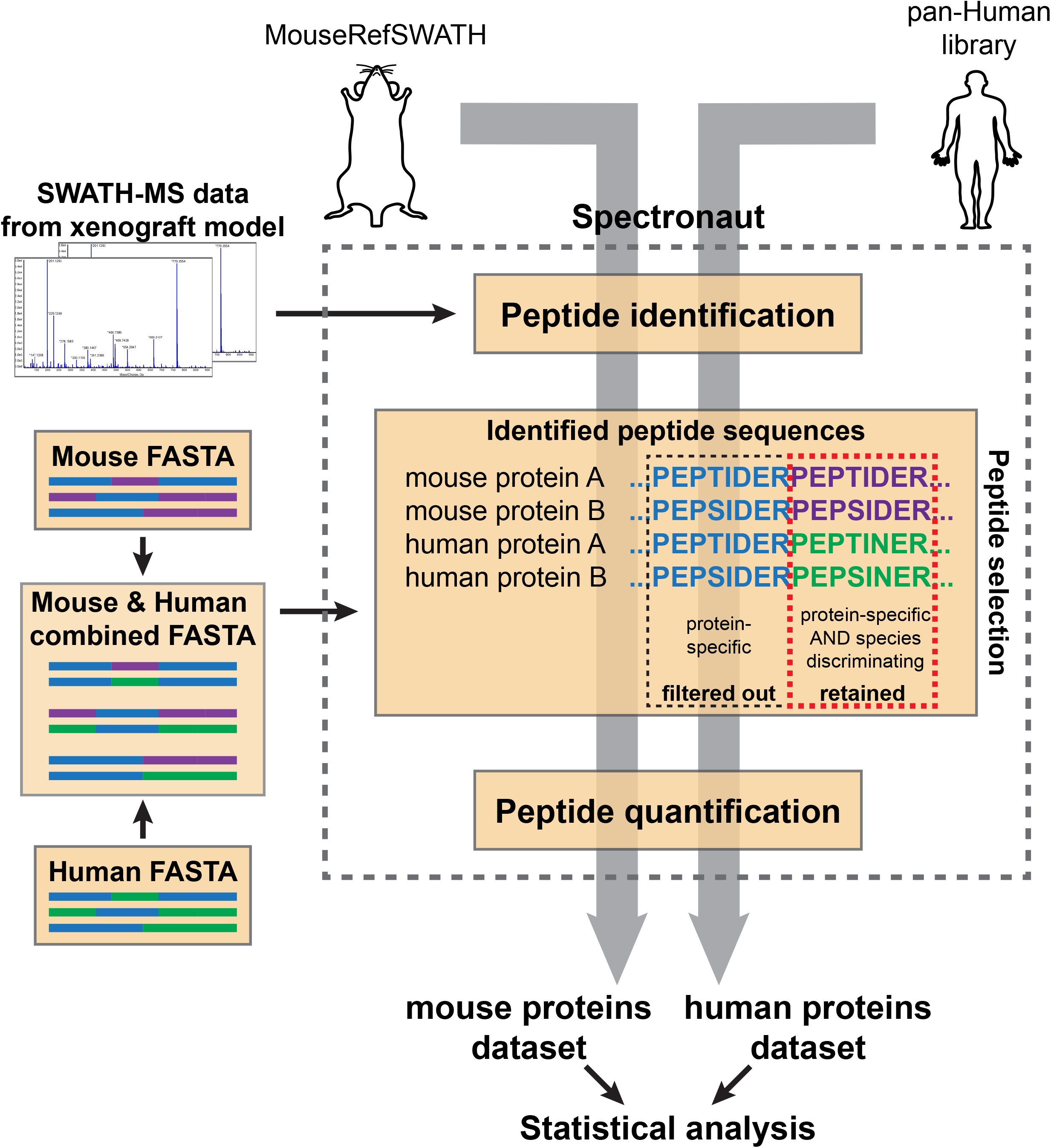
XenoSWATH workflow of the species-specific deconvolution analysis pipeline. The ‘bulk tumour xenograft’ acquired SWATH-MS data is first subjected to two separate searches by either the MouseRefSWATH or the pan-Human library in the Spectronaut software to identify protein-specific (proteotypic) peptides. A combined FASTA file of human and mouse *in silico* digested peptides is then generated to enable subsequent peptide quantification of species-discriminating proteotypic peptides. For peptide quantification, Spectronaut will compare the sequences of the identified peptides from either the MouseRefSWATH or pan-Human library searches with the peptides sequences in the combined FASTA file. Because protein-specific (proteotypic) peptides that are not species-discriminating (blue) occur more than once in the combined FASTA file, Spectronaut filters these peptides out. Only peptides which are both protein-specific and species-discriminating (purple for mouse and green for human) in the combined FASTA file are retained and subjected to quantification. The output of this pipeline is two quantified datasets, one specific for mouse proteins and the other for human proteins.

### SWATH-MS analysis and deconvolution of tumour (human) and host mammary gland (mouse) proteomic alterations in a xenograft model of DCIS progression

We applied the XenoSWATH deconvolution pipeline to quantify the species-specific proteomic alterations associated with DCIS progression to IBC in an orthotopic mouse intraductal breast DCIS (MIND) xenograft model. The MIND model is based on the injection of human breast DCIS cells such as the MCF10DCIS.com cell line into the mouse mammary duct (Figure 4A) [35–37]. Compared to the other breast cancer xenograft models such as mammary fat pad injection, the MIND model has been shown to better recapitulate the mammary gland microenvironment [36], a key regulator of breast cancer progression [31, 38], and is therefore a more clinically relevant model for this disease. In our experiments, MCF10DCIS.com-Luc cells were injected intraductally into the mammary glands of mice where they form tumours that faithfully model the process of DCIS progression over the course of 10 weeks (Figure 4B-D) [35]. Tumours at 4 weeks post injection (referenced as 4w, number of biological replicates n=7) mimic non-invasive DCIS lesions, while after 6 weeks of growth (6w, n=7), tumours start microinvading into the surrounding tissue and finally progress to full IBC at 10 weeks (10w, n=8) post injection (Figure 4D). There is a significant increase in size of the 10w lesions compared to the 4w (p=0.0012) and 6w (p=0.0059) lesions but no significant difference in tumour size between 4w and 6w lesions (p=0.317) (Figure 4B-C) when most lesions at 6w remain within the duct and only a few cells are microinvading out of the ducts.

**Figure 4.**
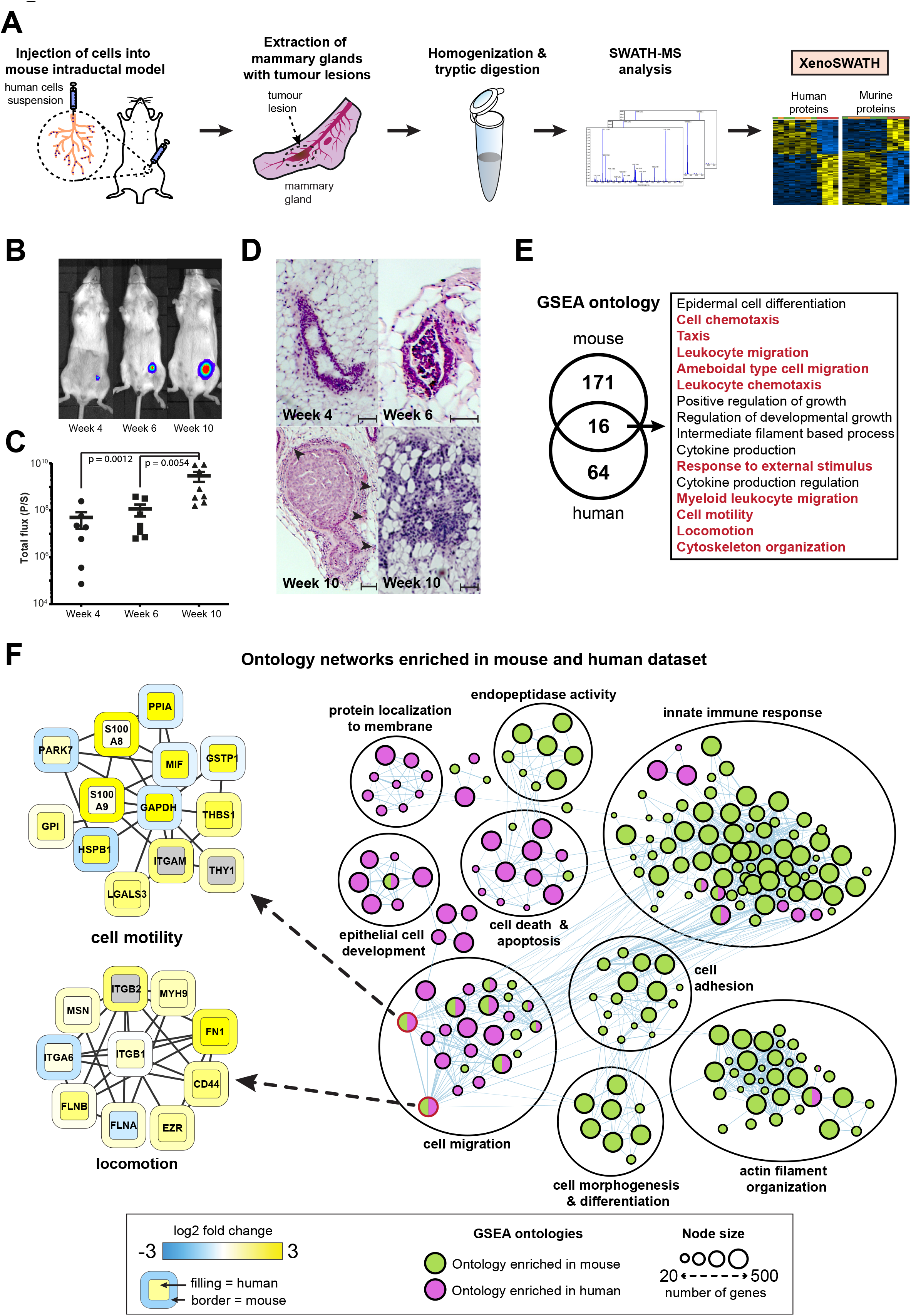
Quantitative proteomic profiling of DCIS progression to IBC in the mouse intraductal (MIND) xenograft model. A) Experimental workflow of SWATH-MS analysis of tumour xenografts. MCF10DCIS.com-Luc cells were injected orthotopically into mouse mammary gland ducts through the nipple. Whole mammary glands with tumours were removed at 4, 6 and 10 weeks (w) post-injection and subjected to sample preparation prior to SWATH-MS data acquisition and analysis by the XenoSWATH pipeline. B) Representative bioluminescence images as measured by IVIS at 4, 6 and 10w post-injection. C) Total bioluminescence flux reflecting tumour size in MIND model at 4, 6 and 10w post-injection (n=7-8). p-value represents statistical significance of the difference between the sample groups calculated by two-tailed Mann-Whitney test. D) H&E images showing initial DCIS lesion formation in the MIND model at 4w which progresses to form an invasive tumour at 10w post-injection. Tumour microinvasion in lesion 10w post-injection is indicated by arrowheads. E) Venn diagram showing the overlap of enriched ontologies for proteins that are significantly upregulated in the 10w specimens versus the 4 and 6w specimens as identified by GSEA with FDR<0.1. Ontologies associated with cell motility, migration and cytoskeleton organization are highlighted in red. F) Network depicting ontologies enriched in human (purple) and mouse (green) datasets (10w versus 4w and 6w) and their overlap as identified by GSEA. Detailed protein networks for two selected ontologies (cell motility and locomotion) are shown. In these networks, the protein node border colour represents the mouse dataset while the protein node fill colour represents the human dataset. Proteins that are not detected are represented by grey.

Whole mammary glands with tumour (n=4) were collected after 4, 6 and 10 weeks post injection and subjected to SWATH-MS analysis (Figure 4A). When processed through the XenoSWATH pipeline (Figure 3), we successfully quantified 2,086 murine and 1,177 human proteins in all samples across the three time-points (Table S1). Our analysis showed no significant differences in protein expression levels in both human and murine datasets between the 4w and 6w specimens, indicating that the non-invasive DCIS lesions (4w) and tumours with microinvasion (6w) harbour similar proteomic profiles at this level of resolution. Comparing the 4w and 6w data with the 10w lesions led to the identification of 327 murine and 247 human proteins that showed significant changes in protein expression (Table S2).

Gene set enrichment analysis (GSEA) was performed to identify functional ontologies that were enriched in the murine and human proteomic datasets upon DCIS progression to IBC (Figure 4E and Table S3). This analysis showed a number of ontology networks that were almost exclusively enriched in either in the human tumour cells (cell death & apoptosis, protein localisation to membrane and epithelial cell development) or the mouse stroma (cell adhesion, actin filament organisation and innate immune response) (Figure 4F). Notably, we identified 16 overlapping ontologies that were upregulated in both species, the majority of which were associated with increased cell migration (Figure 4E-F). An assessment of the upregulated proteins in a subset of these overlapping ontologies revealed a complex regulation of protein networks in both the human and mouse compartments. For instance, within the networks regulating cell motility and locomotion (Figure 4F), specific proteins were upregulated in the tumour cells only [e.g macrophage inhibitory factor (MIF) and peptidylprolyl isomerase A (PP1A)], host cells only [e.g. S100a8 and S100a9]; or in both compartments [e.g. Galectin-3 (LGALS3), Fibronectin (FN1), Ezrin (EZR), CD44 antigen (CD44)]. Employing the deconvolution pipeline, our analyses demonstrates for the first time that the temporal upregulation of cell migration pathways consistent with DCIS to IBC progression is not restricted only to the tumour cells but is found to also operate in the host tumour microenvironment. This proof-of-principle experiment demonstrates the utility of our XenoSWATH deconvolution pipeline for the comprehensive mapping of temporal alterations in the tumour versus host proteome in tumour xenograft models and its ability to capture relevant tumour biology for subsequent investigation.

## Discussion

In this study, we have built the first comprehensive mouse reference spectral library (MouseRefSWATH) for SWATH-MS applications. The MouseRefSWATH library is generated from proteomic datasets acquired across wide range of murine tissue samples, primary cells and cell lines which facilitates versatile use in proteomic profiling of various sample types. We demonstrate its utility in two publicly available SWATH-MS datasets where the use of MouseRefSWATH identified and accurately quantified >80% of the proteins compared to study-specific libraries with a Pearson’s correlation coefficient for protein quantification of at least 0.78. In both datasets, the MouseRefSWATH library further identified new proteins which would have otherwise not been possible with study-specific libraries. In addition to removing the need to generate study-specific libraries for future murine SWATH-MS experiments, the benefits of utilising the larger MouseRefSWATH library include the fact that individual proteins are represented by a higher number of precursor ions resulting in better accuracy and higher confidence in protein quantification.

We have also extended the use of the MouseRefSWATH library and developed a novel analysis pipeline called XenoSWATH that enables deconvolution of murine and human proteins from ‘bulk tumour’ xenograft proteomic measurements through the identification of species-discriminating proteotypic peptides. The lack of tools to perform a deep analysis of tumour (human) and host (mouse) molecular alterations *in situ* have limited our ability to study the role of the tumour microenvironment in driving tumour progression. *In silico* approaches have been developed to deconvolute mouse and human reads in next generation DNA and RNA sequencing data derived from tumour xenografts but such approaches do not yet exist for proteomic data analysis [39]. The only published application of SWATH-MS in a xenograft model thus far has focused on measuring the murine stromal component by pre-enrichment of mouse cells with immunoaffinity chromatography while completely omitting the human tumour cells [33]. XenoSWATH presents a paradigm shift in the ability to distinguish and comprehensively map proteomic alterations in both the human and mouse compartments and provides a new approach to investigating tumour cell-microenvironmental interactions in cancer initiation, progression and therapy response. With the recent development of human tumour xenograft models in immunodeficient zebrafish [40], we anticipate that XenoSWATH can be readily extended to the study of host and tumour cell responses in these models by similarly utilising the zebrafish reference spectral library available in SWATHAtlas [17].

As proof-of-principle, we undertook a quantitative proteomic analysis in the MIND model and provide the first characterisation of host stroma and tumour cell proteomic alterations that occur during DCIS to IBC progression. Our analysis reveal complex network alterations in both compartments, and while the human tumour cells show an expected upregulation of migration pathways during progression to invasive disease, we make the unanticipated discovery that the mouse stroma also displays an enrichment of cell migration and motility networks upon DCIS progression. In the mouse dataset, the highest observed increase in migration-associated protein expression levels between early (4w & 6w) versus late (10w) timepoints were in the S100a8 and S100a9 proteins (Table S2). These two proteins together form the calprotectin complex and stromal cells secreting calprotectin have been previously observed in the microenvironment of pancreatic cancer [41]. Additionally S100a8 has been reported to increase the migration and proliferation of colorectal and pancreatic cancer cells *in vitro* [41] and has also been associated with metastasis formation in breast cancer [42]. MIF was found to be exclusively upregulated in the human DCIS.com cells at 10 week post-injection but not in the mouse stromal cells. Increased levels of MIF has been reported in many cancer types including pancreatic cancer [43], hepatocellular cancer [44] or head and neck cancer [45]. Experiments with murine breast cancer cell line 4T1 have shown that overexpression of MIF promotes tumour metastasis [46] and protects cancer cells from immunogenic cell death [47]. Finally, we determine that LGALS3, CD44, FN1 and EZR were upregulated in both tumour and host datasets. These proteins have been extensively shown in published literature to regulate cytoskeleton remodelling, epithelial-mesenchymal transition and increased cell motility [48–50], all of which are important in breast cancer progression to invasive disease. These examples highlight the utility of this deconvolution pipeline in dissecting the individual roles of the tumour cells and the associated stroma in cancer biology. This dataset serves as a rich resource of tumour and microenvironmental proteins for future functional investigation, candidate biomarkers for stratifying DCIS patients with increased risk of progressing to IBC, as well as targets for drug discovery for delaying DCIS progression.

There are several limitations to the XenoSWATH pipeline. Because the method focuses on quantifying peptides that are both proteotypic and species-discriminating, a significant number of proteins with high sequence similarity between mouse and human are filtered out and lost during the process, resulting in the quantification of a reduced subset of the proteome. Despite this limitation, we are still able to readily identify key pathways that are operating in both the human and mouse compartment. Given that the stroma is composed of a number of different cell types including fibroblasts, endothelial cells and immune cells; as with other population-level based measurement methodologies, the XenoSWATH pipeline is unable to resolve the individual contribution of specific stromal cell types to the aggregate proteomic data. Extending *in silico* transcriptomic deconvolution strategies that are currently being used to estimate stromal cell types to proteomic data may provide better resolution and address this shortcoming in future [51].

Collectively, our study has generated the MouseRefSWATH comprehensive mouse reference spectral library as a standardised community resource for use in future mouse SWATH-MS studies, which will not only remove the need for the generation of study-specific libraries but will also improve inter-laboratory reproducibility. We further present a new XenoSWATH analysis pipeline for species-specific deconvolution of xenograft proteomic data which opens new possibilities for the in-depth assessment of tumour-host interactions in murine xenograft models. Moving forward, we anticipate that these tools will have broad applications in addressing key biological research questions involving this widely used model organism.

## Methods

### Murine organs, primary cells and cell lines

Isolation of T-cells was carried out under Swiss animal experiment regulations. All other animal work was carried out under UK Home Office project and personal licenses following local ethical approval from the Institutional Animal Ethics Committee Review Board and in accordance with local and national guidelines. Mammary gland, liver and lung organs were dissected from 14-week-old to 18-week-old virgin female SCID Beige mice. Brain, heart and kidney organs were dissected from 6 months old female NCR nude mice. Lymph nodes were dissected from 2-4 months old C57BL/6N male and female mice. All tissue specimens were briefly washed in cold phosphate-buffered saline (PBS) to remove excess blood and immediately snap frozen in liquid nitrogen and stored at −80 °C.

Primary CD8+ effector T-cells were isolated from splenocytes of OT-I mice and cultured in Roswell Park Memorial Institute (RPMI) medium with 10% foetal bovine serum (FBS) (Gibco), 1% penicillin-streptomycin and 0.1% β-mercaptoethanol. In order to activate OT-I CD8^+^ T-cells, OT-I splenocytes were treated with 1 μg/ml OVA257-264 peptides in the presence of 10 ng/ml IL-2 for 3 days while the non-stimulated resting CD8^+^ T-cells were collected after culturing with IL-2 alone. Immortalized normal (NF1) and cancer-associated fibroblasts (CAF1) [24] were cultured in Dulbecco’s Modified Eagle Medium (DMEM) supplemented with 10% FBS, 1x GlutaMAX (Gibco), 0.5% penicillin-streptomycin and 1x insulin-transferrin-selenium A (ITS-A) (Gibco). NIH-3T3 cells were cultured in DMEM supplemented with 10% FBS/100units/ml penicillin/100mg/ml streptomycin. C2C12 and 4T1 cells were cultured in RPMI supplemented with 10% FBS/100units/ml penicillin/100mg/ml streptomycin. Ba/F3 cells were cultured in the same media as 4T1 cells with the addition of 5ng/ml IL-3. All cells were cultured in 95% air/5% CO2 atmosphere at 37°C.

### Tissue and cell sample processing

Tissue samples were cut into small pieces and placed into precooled tubes. High salt homogenization buffer consisting of 50mM Tris-HCl (pH 7.4), 0.25% 3-[(3-Cholamidopropyl)dimethylammonio]-1-propanesulfonate hydrate (CHAPS, Sigma), 25mM EDTA, 3M NaCl (Sigma) and 10 KIU/ml aprotinin was added at 4 ml/g of tissue and samples were homogenized by 2 × 30 s pulses on ice with a LabGEN125 (Cole-Parmer) homogenizer. Homogenized samples were rotated for 20 min at 4 °C. Proteins in homogenate were acetone precipitated by mixing with 4 volumes of ice cold acetone, vortexed and incubated 2 h at −20 °C. Samples were spun for 15 min at 15,000 rpm, 4 °C, supernatant removed and resulting pellet was resuspended in 0.3 ml of urea buffer consisting of 8M urea (Sigma), 100 mM ammonium bicarbonate (Sigma). Protein concentration in samples was measured using the Pierce 660nm Protein assay (Thermo Scientific) as per manufacturer’s instructions and stored at −80 °C until further processing.

Cells were harvested and lysed in 8 M urea buffer and protein concentration was measured by Pierce BCA assay (Thermo Scientific) as per manufacturer’s instructions. Cell lysates were stored at −80 °C until further processing

For each sample lysate, 200 μg of total protein was reduced with 20mM dithiothreitol (Sigma) at 56 °C for 40 min and alkylated by 30mM iodoacetamide (Sigma) at room temperature for 25 min in the dark. Samples were diluted to a final concentration of 2M urea, 100mM ammonium bicarbonate and digested at 37 °C overnight with 4 μg of sequencing grade trypsin (Promega). Digestion was stopped by acidification to pH<4 with trifluoracetic acid (Sigma) and resulting peptides were desalted on SepPak C18light (Waters) cartridges as per manufacturer’s instructions. Desalted peptides were dried in a SpeedVac concentrator and stored at −20 °C until fractionation.

### Fractionation by strong cation exchange (SCX) chromatography

Dried peptides were resuspended in 100 μl of buffer SCX-A consisting of 10mM NH_4_COOH (Sigma) in 20% acetonitrile (Thermo Fisher), pH 2.7, vortexed, sonicated for 5 min and spun at 15,000 rpm for 1 min. The supernatant was loaded on a 2.1 × 100 mm Polysulfoethyl A column with 5 μm, 200 Å particles (PolyLC Inc.) and eluted with buffer SCX-B (500mM NH_4_COOH in 20% acetonitrile, pH 2.7) using a gradient of 0-10% of B in 2.5 min, 10-50% of B in 20 min, 50-100% of B in 7.5 min and 100% of B for 10 min. Twelve fractions were manually collected over 39 min with fraction 1 collected from 0 to 12 min and fraction 12 from 32 to 39 min. The remaining 10 fractions were collected at 2 min intervals between 12 to 32 min. All SCX fractions were dried in a SpeedVac concentrator.

### Fractionation by high pH reverse phase (HpH RP) chromatography

Dried peptides were resuspended in 100 μl of buffer HpH-A (0.1% NH_4_OH), vortexed, sonicated for 5 min and spun at 15,000 rpm for 1 min. The supernatant was loaded on a 2.1 × 150 mm XBridge BEH C18 column with 5 μm, 130 Å particles (Waters) and eluted with buffer HpH-B (0.1% NH_4_OH in acetonitrile) using a linear gradient of 0-50% B over 60 min. Fractions were automatically collected between 5 and 50 min every 30 s into 96 well plate and fractions were pooled into 12 fraction (columns pooled) for 3T3 cells or 8 fractions (rows pooled) for 4T1, BaF3 and C2C12 cells. Pooled fractions were dried in a SpeedVac concentrator.

### DDA MS data acquisition

All fractions were resuspended in 20 μl of buffer A (2% acetonitrile, 0.1% formic acid) and peptide concentration was measured using the 280 nm NanoDrop assay. 1 μg of total peptide was analysed in DDA mode on an Agilent 1260 HPLC coupled to a TripleTOF 5600+ mass spectrometer equipped with NanoSource III. Each fraction was spiked with 0.1 μl of iRT calibration mix (Biognosys, AG) and loaded onto a 0.3 × 5 mm ZORBAX C18 (Agilent Technologies) trap column. Peptides were separated on a 75 μm × 15 cm analytical column packed with Reprosil Pur C18AQ beads, 3 μm, 120 Å (Dr. Maisch, GmbH) with a manually pulled integrated spraying tip. A linear gradient of 2-40% of buffer B (98% acetonitrile, 0.1% formic acid) in 90 min and flow rate of 250 nl/min was used for peptide separation. All data were measured in positive mode. Full profile MS scans were acquired in the m/z mass range of 340-1500 with 250 ms filling time, MS/MS scans for the 20 most intense ions with charge state from 2+ to 5+ were acquired in m/z mass range of 280-1500 with 100ms filling time. Dynamic exclusion of fragmented ions was set to 12 s.

### DDA MS data processing and MouseRefSWATH reference spectral library generation

All acquired DDA datasets were searched by SpectroMine (ver 1.0.21621.7 Sapphire) software (Biognosys AG) against a Swissprot mouse database (downloaded on 26/10/2018) with added iRT peptide sequences. Carbamidomethylation of cysteines was set as a fixed modification, oxidation of methionine and proline, deamidation of glutamine and asparagine, acetylation of protein N-terminus were set as variable modifications. A maximum of 2 missed cleavage sites were allowed during the search. False discovery threshold on peptide and protein level was set to 1% to filter search results.

To generate final reference spectral library, retention times in each run were individually calibrated using iRT calibration peptides. Runs with low result of calibration fit (R^2^<0.8) were removed. To avoid inflation of FDR during the library generation from multiple datasets, Search Archives of all datasets were combined in SpectroMine and the MouseRefSWATH reference spectral library was built using following the parameters: minimum 3 and maximum 6 transitions for each precursor, library-wide protein and peptide FDR threshold of 1%. The MouseRefSWATH libraries is deposited in ProteomeXchange with identifier PXD017209.

### Comparative analysis of publically available SWATH-MS data

Publicly available datasets (PXD006382, PXD005044) were downloaded from data repository via the ProteomeCentral portal [52] and analysed by Spectronaut (version 13.6.190905) software (Biognosys AG). Either the MouseRefSWATH library or the relevant study-specific spectral library was uploaded into Spectronaut and all samples were processed using FDR threshold of 1% on peptide and protein levels. Mitochondrial proteins were identified by matching the list of quantified proteins with the MitoCarta 2.0 database [53]. It should be noted in the original published studies, older versions of the Spectronaut software with different settings were used by the authors which leads to the discrepancies in number of reported quantified proteins in our study when compared to the original published data. Detailed Spectronaut settings for this study are shown in Supplementary methods

### Mouse intraductal DCIS (MIND) model

The MIND model was utilised in our studies as previously described [35, 54]. Briefly, a suspension of 5 × 10^4^ MCF10DCIS.com-Luc cells was injected intraductally into mammary gland ducts of 6-10 week-old SCID-beige female mouse (n=7-8). Luminescence of the lesions was measured by *in-vivo* imaging assay (IVIS) on IVIS Illumina II (Perkin Elmer) to monitor tumour growth. Whole mammary glands with tumour were collected after 4, 6 or 10 weeks post injection, washed in cold PBS and freshly frozen. For each condition, 4 biological replicates were further processed and analysed by SWATH-MS. Haematoxylin and eosin (H&E) staining was performed on formalin-fixed and paraffin embedded samples and tissue images were scanned on Nanozoomer XR (Hamamatsu Photonics) automated slide scanner.

### MIND model sample processing and SWATH-MS data acquisition

Whole mammary glands with tumour cells were homogenized in high salt homogenization buffer, proteins were precipitated with ice-cold acetone at −20 °C, centrifuged and the resulting pellet was resuspended in 0.3 ml of urea buffer. 20μg of total protein was digested in solution by trypsin as described above, desalted on OMIX tips as per manufacturer’s instructions and dried in a SpeedVac concentrator. Dried samples were resuspended in 20μl of buffer A and analysed in SWATH-MS mode on an Agilent HPLC coupled to TripleTOF 5600+ mass spectrometer. 1 μg of sample was spiked with 0.1 μl of iRT peptides and loaded onto a 0.3 × 5 mm ZORBAX C18 (Agilent Technologies) trap column. Peptides were separated on a 75 μm × 15 cm analytical column packed with Reprosil Pur C18AQ beads, 3 μm, 120 Å (Dr. Maisch, GmbH) with a manually pulled integrated spraying tip. A linear gradient of 2-40% of buffer B in 120 min and flow rate of 250 nl/min was used for peptide separation. All data were acquired in positive ion mode in SWATH mode using cycles comprising of one 100ms profile MS scan over the m/z mass range of 340-1500, followed by 60 SWATH fragmentation windows with a fixed width of 12 Da over the m/z range of 380-1100 and filling time of 50 ms. The SWATH-MS data is deposited in ProteomeXchange with identifier PXD017209.

### Implementing XenoSWATH species specific deconvolution pipeline and MIND model SWATH-MS data processing

A combined FASTA file was generated in NotePad text editor from individual FASTA files containing human (20,316 protein sequences), mouse (16,997 protein sequences) and iRT peptides. The human and mouse FASTA files were downloaded from SwissProt (downloaded on 26/10/2018) and the FASTA file with iRT sequences was downloaded from Biognosys website (https://www.biognosys.com/shop/irt-kit#SupportMaterials). The acquired MIND model SWATH-MS data were processed in Spectronaut using the MouseRefSWATH reference spectral library, the published pan-Human reference spectral library [15] and the combined FASTA file. The SWATH-MS data was first searched using the MouseRefSWATH library. Using the combined FASTA file, Spectronaut selects only the mouse species discriminating proteotypic peptides from the MouseRefSWATH library for the quantification of murine proteins. The same approach was repeated with pan-Human library for the quantification of human species discriminating proteotypic peptides. All searches were performed with 1% FDR threshold on peptide and protein level (detailed Spectronaut settings are shown in Supplementary methods). In this manner, two separate proteomic datasets were obtained, one for the tumour component (human) and the other for the host stromal compartment (mouse). The datasets were separately quantile normalized using proBatch package [55] in R and statistically analysed by ANOVA in Perseus [56]. Analysis of ontologies enriched in 10w versus 4w & 6w was performed using GSEA desktop application against human or mouse MSigDb of biological processes [57, 58]. FDR threshold was set to 0.1 using genotype permutations due to low number of biological replicates. Protein networks were visualized in Cytoscape (version 3.7.1) [59] and clustered by the MCL clustering algorithm in clusterMaker2 Cytoscape plugin[60].

## Acknowledgements

P.H.H is supported by grants from the Institute of Cancer Research (ICR), Breast Cancer Now (2013NovPhD185 and 2014NovPR360) and Cancer Research UK (C36478/A19281). B.A.H and R.C.N are supported by Breast Cancer Now programmatic funding. S.E.A is supported by grants from Cancer Research UK (CRUK/A19763) and Medical Research Council (MC_U12266B). F.C. is supported by grants from the Institute of Cancer Research, the Ramon y Cajal Research Program (RYC-2016-20352; FSE/AEI), and MCIU/AEI/FEDER (RTI2018-096778-A-I00). P.-C.H. is supported in part by the SNSF project grants (31003A_182470) and the European Research Council Staring Grant (802773-MitoGuide).

## Supplementary tables

**Table S1:** Full proteomic dataset of human and mouse proteins obtained from MIND model experiment as processed by XenoSWATH deconvolution pipeline. Confidence of the protein identification is reported as FDR value.

**Table S2:** List of mouse and human proteins with significantly altered expression in 10w samples compared to 4w & 6w samples. Log2 fold change for each protein is reported along with the t-test FDR value.

**Table S3:** GSEA normalized enrichment scores and FDR values for mouse and human overlapping ontologies based on proteins which are upregulated in 10w versus 4w & 6w samples (Figure 4E).

